# Whole organ volumetric sensing Ultrasound Localization Microscopy for characterization of kidney structure

**DOI:** 10.1101/2023.08.31.555780

**Authors:** Georges Chabouh, Louise Denis, Sylvain Bodard, Franck Lager, Gilles Renault, Arthur Chavignon, Olivier Couture

## Abstract

Glomeruli are the filtration units of the kidney and their function relies heavily on their microcirculation. Despite its obvious diagnostic importance, an accurate estimation of blood flow in the capillary bundle within glomeruli defies the resolution of conventional imaging modalities. Ultrasound Localization Microscopy (ULM) has demonstrated its ability to image in-vivo deep organs in the body. Recently, the concept of sensing ULM or sULM was introduced to classify individual microbubble behavior based on the expected physiological conditions at the micrometric scale. In the kidney of both rats and humans, it revealed glomerular structures in 2D but was severely limited by planar projection. In this work, we aim to extend sULM in 3D to image the whole organ and in order to perform an accurate characterization of the entire kidney structure. The extension of sULM into the 3D domain allows better localization and more robust tracking. The 3D metrics of velocity and pathway angular shift made glomerular mask possible. This approach facilitated the quantification of glomerular physiological parameter such as an interior traveled distance of approximately 7.5 ± 0.6 microns within the glomerulus. This study introduces a technique that characterize the complete kidney phisiology which can serve as a method to facilite pathology assessment. Furthermore, its potential for clinical relevance could serve as a bridge between research and practical application, leading to innovative diagnostics and improved patient care.

## Introduction

Glomeruli represent the filtration units of the kidney. In clinical practice, the change in the estimated glomerular filtration rate (GFR) - measured by blood or urine tests - is directly related to the degree of renal dysfunction^1^. In fact glomerular diseases represents the highest impact on kidney allograft loss^28^. In contrast to laboratory measurements, imaging enables direct visualization and quantification of kidney tissue remodeling, inflammation, and fibrosis^2^. However, accurately estimating blood flow at the scale of tenths of a millimeter within glomeruli defies the resolution capabilities of conventional ultrasound, computed tomography (CT), and magnetic resonance imaging (MRI)^3^.

Ultrasound Localization Microscopy (ULM) has demonstrated its capacity to image in-vivo deep organs in the body^4–6^, including the human renal graft^7^. It has emerged as a powerful technique that has successfully overcome the limitations imposed by the diffraction barrier, enabling the acquisition of high-resolution images in-vivo. The technology has demonstrated its potential for imaging the microcirculation in various organs such as brain for various organs such as brain^29^, spinal cord^30^ and kidney^31,38^ in rodents. However, to fully realize the clinical benefits of ULM, there remains a crucial gap that needs to be addressed, specifically in integrating ULM into routine medical practice and define new biomarkers. Beyond the mere attainment of exquisite images, there exists the pressing need to explore and exploit ULM’s capabilities for advanced medical analyses, such as the estimation of blood volume and other relevant clinical features as discussed in a recent review^32^. Therefore, the current focus should shift towards harnessing ULM’s potential as a versatile tool for quantitative and functional assessments, paving the way for its practical implementation in clinical settings. Various research studies goes into this direction such as using the vascular properties to study obesity through the kidney^20^, or for functional^33^ and dynamical imaging^34^ of the brain.

In a recent study^8^, we introduced the concept of sensing Ultrasound Localization Microscopy (sULM) in 2D, revealing the kidney’s functional structures (glomeruli) in both rats and humans grafts. By dividing the data into two categories of velocity: slow microbubbles in the medulla and glomeruli and fast ones in the bigger arteries, all kidney regions were characterized using specific metrics relying on the kinetics of these microbubbles.

However, a significant constraint in studying intrinsic microbubble behavior stemmed from the presence of vessels running transverse to the imaging plane. As the microbubble deviates from the 2D imaging plane, its intensity progressively diminishes until it vanishes. This hinders the continuous tracking of the microbubble throughout its intravascular course. Moreover, relying on a lone 2D imaging plane restricts the potential diagnostic capabilities of sULM. While a conceivable solution involves multi-plane imaging, although acquisition time becomes important^35^. Conversely, accurate quantification are hindered through 2D projections. Thus, a comprehensive solution lies in whole-organ 3D imaging, which not only addresses the aforementioned issues but also reduces reliance on user interpretation.

In a recent work, 3D transcranial ULM was proposed as a discriminator between ischemic and hemorrhagic stroke in early phase for a preclinical model^18^. Various studies, propose a shift from qualitative imaging methods to a more quantitative approach. In fact, 3D ULM was also used to study vasodilatation^36^ in the brain as well as in the heart^37^ both in pre-clinical studies.

In the presented work, our objective is to demonstrate that the implementation of 3D sULM enables accurate tracking of microbubble pathways throughout the entire intravascular trajectory. Additionally, we emphasize the capability of 3D sULM to achieve precise quantification of various physiological parameters, including factors like glomerular blood flow and size, working towards a more accurate estimation of the glomerular filtration rate (eGFR).

## Materials and Methods

### In-vivo rat model

All animal experiments were performed in accordance with the ARRIVE guidelines and approved by the local ethics committee (ethics committee on animal experimentation n^*circ*^034). The protocol is registered by the ministry of research under number #33913-2021082311153607. In accordance with the 3R rules, the number of animals in our study was kept to the necessary minimum. Experiments were performed on 8 Wistar male rats (Janvier Labs, Charles River, Envigo) aged between 8 and 12 weeks. Only 5 rats were included due to technical problem with data acquisitions and saving. All 8 rats were used to set-up and validate the micro-CT terminal perfusion of baryum sulfate solution. Also, they allowed the realization of a complete 3D sULM mapping. Rats were anesthetized during the entire experiment (4% Isoflurane) and maintained at a temperature of 37°C using a heating table. The animals died during the injection of the micro-CT contrast medium detailed later in this section.

### Ultrasound acquisitions

To perform the ultrasound acquisition, the left kidney was externalized through an incision in the abdomen (See Fig.2 **a**). The organ was then placed on an absorber and fixed with a needle to avoid movement. A 25G catheter was placed in the tail vein to perform microbubbles injections (SonoVue, Bracco, Italy) needed for volumetric sULM. The animal was under isoflurane anesthesia (4% for induction, 2.5% for maintenance) with appropriate analgesia (subcutaneous injection of 0.1 mg/Kg of buprenorphine 30 minutes prior to experiments).

Ultrasound acquisitions were then performed with a 256-channels research ultrasound scanner (verasonics, Kirkland, USA) and an 8 MHz multiplexed matrix probe (Vermon, France). Five hundred blocks were acquired, with 200 images per block and a framerate of 130Hz. Each image being the result of a compounding of 5 plane waves oriented according to ± 5° (in the elevation axis and in the lateral axis). The sequence was decomposed according to a *light* configuration^7^, and lasted for 8 minutes with an injection of 50 *µ*L/min in bolus. The pulse duration is about 2 cycles with a pulse repetition frequency (PRF) of 13.5 kHz. The data were then reconstructed with a classical delay and sum beamforming^16^ on a [98.5, 150, 150] *µ*m grid, before reaching the final sULM resolution of [9.85, 9.85, 9.85] *µ*m^19^.

### Micro computed tomography (micro-CT) acquisitions

Micro-CT data were acquired on the same rats as the ones subjected to ultrasound acquisition at the Live Imaging Platform of the Odontology UFR Montrouge, France (Skyscan 1172, Bruker, [5, 5, 5] *µ*m).

The X-ray opaque contrast medium consists of a mixture of barium sulfate (Micropaque 100 g, Guerbet, France), PBS (Phosphate Buffer Saline, Cochin INSERM U1016), and gelatin. It was injected at the end of the ultrasound acquisitions through an aortic catheterization. All of the animal’s blood was replaced by the contrast medium, i.e., approximately 200 mL/animal. The organ was then stored in a Petri dish for 48 h in 4% of PFA before being scanned in the micro-CT system described above.

Only 4 rats have usable micro-CT data due to the complex nature of the contrast agent injection^6,8^ procedure and the high risk of animal mortality during the process.

No statistical analysis was performed to compare sULM and micro-CT due to non-rigid deformations. Simply the manual registration on the AMIRA (Thermofischer, USA) software, allowed a descriptive assessment in 1 rat, i.e. the rat with the micro-CT and the most complete 3D sULM possible (see Fig.7).

### Whole organ volumetric sULM

The construction pipeline of the 3D sULM mapping is shown in Fig.1. In Fig.1 **a** classical ultrasound Bmode volume reconstruction, **b** clutter filtering using a low-threshold Singular Value Decomposition (SVD) where the first six singular values were annulled, **c** microbubble localization using radial symmetry algorithm^9,10^, and **d** tracking using the Hungarian algorithm^10^. It is worth to mention that in this work, the localization and tracking were done on an unique dataset. Contrary to 2D sULM^6^, where the dataset was divided into two sup-groups: fast moving microbubbles and slow moving microbubbles. Thanks to the volumetric acquisition, we were able to precisely track the microbubble along its entire path within the blood vessels without any loss of visual continuity without the need of separating dataset.

**Figure 1.**
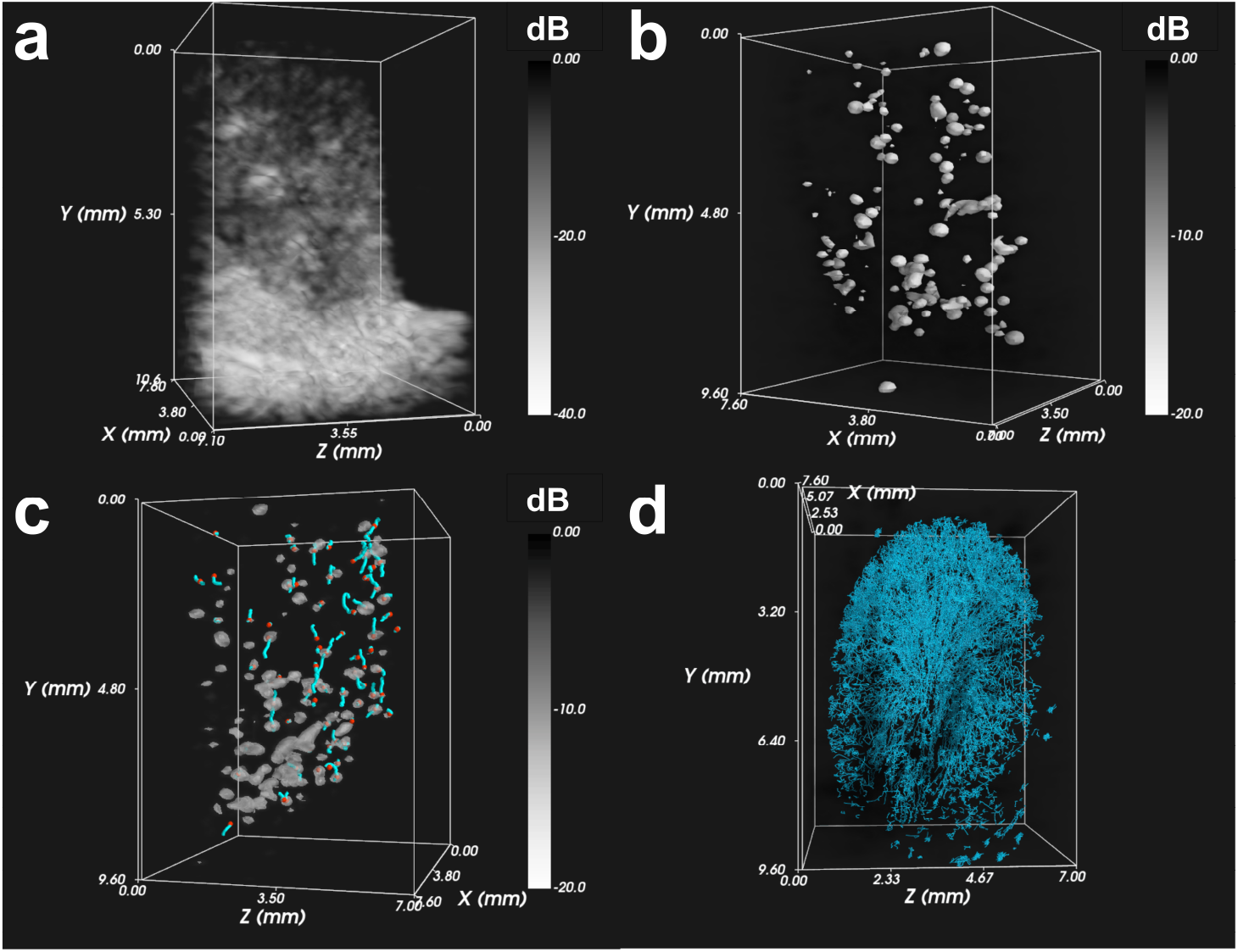
3D sULM steps: **a** B Mode, **b** tissue filtering, **c** microbubble localization and tracking, **d** tracks accumulation.

**Figure 2.**
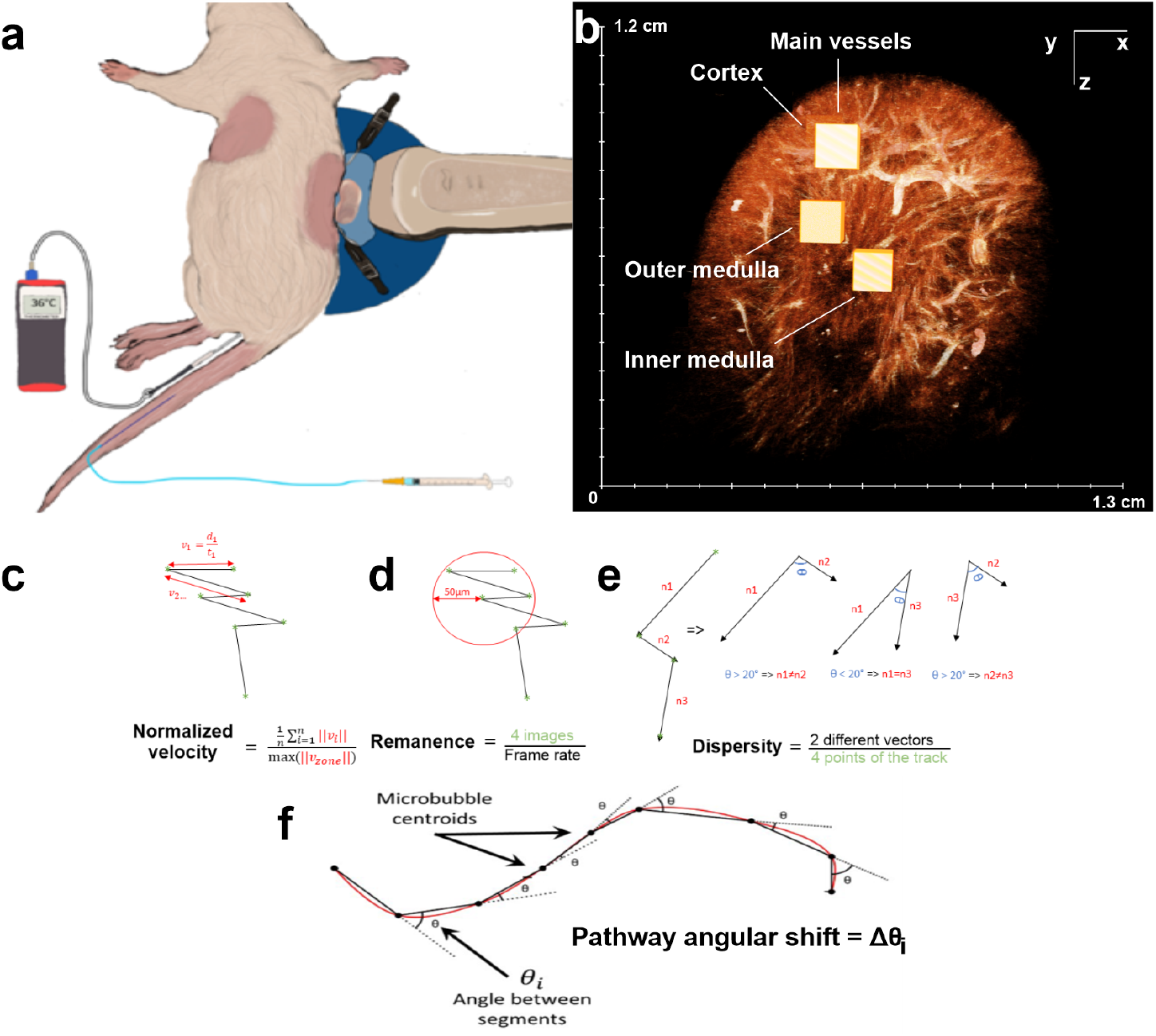
Graphical methods: **a** In vivo setup of the rat kidney **b** 3D ROI of different renal zones were selected from sULM rendering. Representative cubes were chosen for each renal zone on the 3D density mapping and three different metrics used in this study: **c** normalized velocity, **d** Remanance and **e** the pathway angular shift is characterized as the difference between two successive angles, as derived and modified from^17^. Note that PAS does not account for distance normalization between the initial and final points.

### 3D ROI in different renal zones

An arbitrary 3D region of interest (ROI) was delineated within the four principal regions of the kidney: main vessels, inner medulla, outer medulla, and cortex, as depicted in Fig.2 **b**. The ROIs were defined as cubic structures measuring 100 pixels on each side, corresponding to a size of 985 *µ*m. Note that the cortical ROI is not seen from this angle.

### 3D sULM metrics

sULM is built upon an a-priori knowledge of the local environment based on previous invasive microscopy studies such as histology^13^. To highlight specific microbubble behaviour, we we employed two different metrics that we define hereafter.

- Microbubble *velocity*: characterized as the mean displacement magnitude between every two successive points along a trajectory, divided by the corresponding time interval (Fig.2 **c**).
- The *Remanence* Time: established as the maximum period during which a microbubble was tracked within a glomerular sphere, i.e. of 50 *µ*m radius in the rat^13,15^. The center was defined as the median of the list of points constituting the track (Fig.2 **d**).
- The *Dispersity*: corresponded to the number of times a track went in the same direction, by taking the rounded value of the location, with a tolerance of plus or minus 20°, divided by the number of points making up the track (Fig.2 **e**).
- The *Pathway Angular Shift (PAS)*: delineated as the angular difference computed between sets of three consecutive points along a track. To illustrate, envision a track consisting of ten points. The initial trio of points yields two lines, from which an angle can be derived. Subsequently, the subsequent three points generate another pair of lines, yielding a second angle. The disparity between these two angles defines the PAS ((Fig.2 **f**).)

### Statistical analysis

Statistical analyses were performed to evaluate the metrics between the different regions of the kidney using a two-tailed parametric unpaired Student’s t test with GraphPad Prism 9 software. The significance of the results is as follows: *ns* = *P≥* 0.05, ∗= *P ≤*0.05, ∗∗= *P ≤*0.01, ∗∗∗= *P≤* 0.001, ∗∗∗∗= *P≤* 0.0001. All the metrics were calculated inside the above mentioned 3D ROIs as seen in Fig.2 **b** and averaged on 5 different rats (*n* = 5 included).

## 1 Results & Discussion

### 1.1 Vascular mapping along the entire microbubble path

The relatively high volumetric frame rate of our ultrasound acquisitions, enabled us to follow the microbubbles throughout their intravascular journey. Thanks to a single filter and a single tracking algorithm, a complete 3D mapping of the entire kidney in 5 rats was established. We sucessfuly observed the particularities of the microcirculation in each of the regions of the kidney (rat 8 in Fig.3 **a**, and **b**). In the main vessels, the microbubbles move very fast and are subdivided into interlobar arteries, arcuate arteries. The return path has also been reconstructed into an efferent arteriole, arched and interlobar veins (rat 8 in Fig.3 **g** and **h**). In the cortex, some microbubbles ascend in the afferent artery, swirl in the glomerulus before returning to the cortex. We were also able to follow the entire route of the microbubble in the glomerulus (Fig.3 **c** and **d**). Finally, in the medulla, the microbubble path is less trivial but remains a characteristic of the renal physiology. The microbubbles appear to follow a straight path towards the center of the kidney, i.e. along the vasa recta at the level of the Malphigian pyramids (Fig.3 **e** and **f**). After a certain depth inside the inner medulla, a loss of visualisation of these microbubbles is remarked. This can be due to the limitation of the current spatiotemporal filter. In fact, the spatiotemporal based filter employed here might be unable to differentiate fixed or slow moving microbubbles from the tissue signal, since both are highly coherent in space and time. Moreover, the relatively short ultrasound block duration and the time delay between each block of the volumetric ultrasound acquisition present additional explanations for the reduced visualization of slow-moving microbubbles.

**Figure 3.**
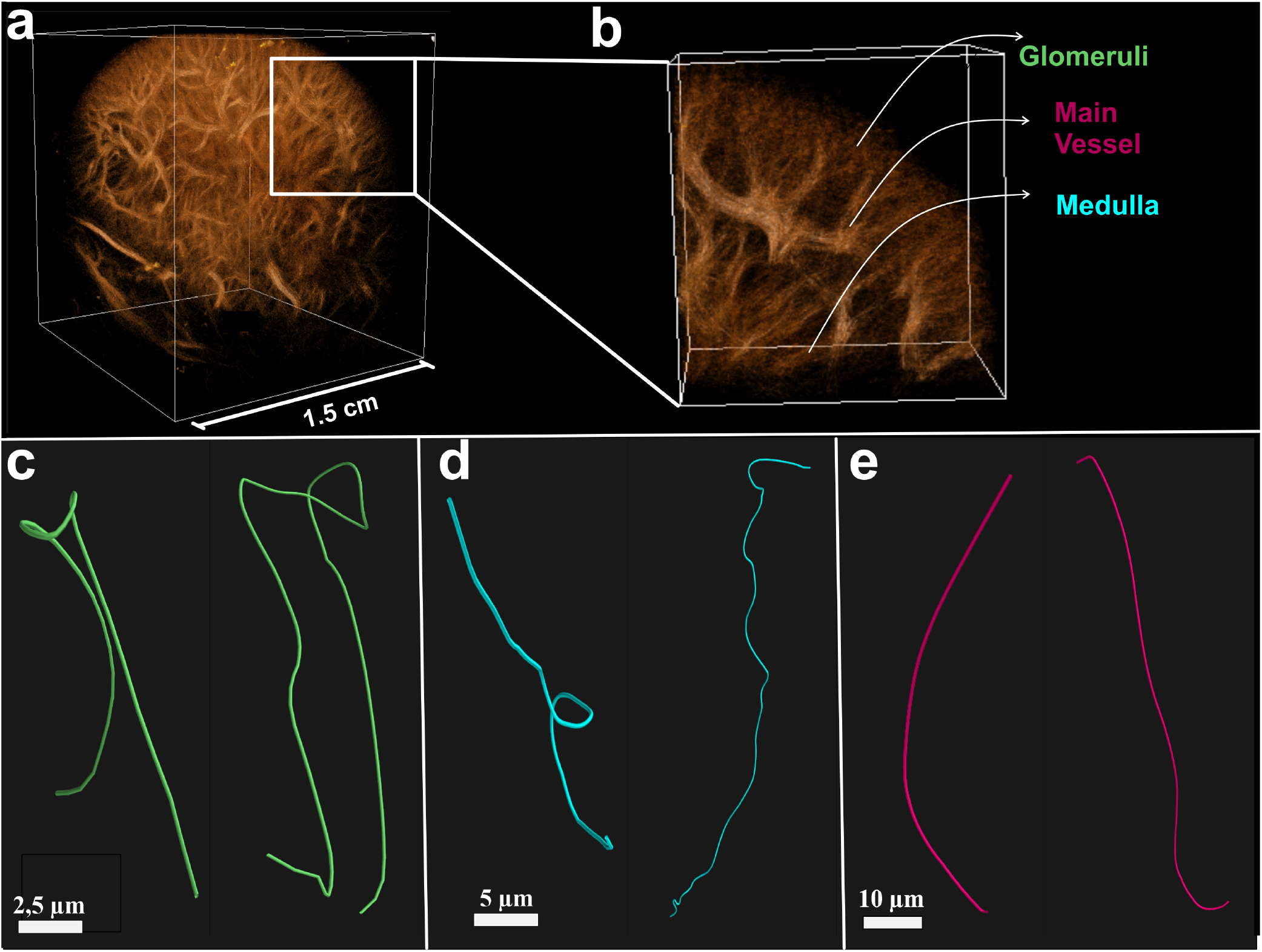
3D ULM mapping: **a** Vascular mapping allows reconstruction of the entire volume of the kidney, **b** Zooming into the cortex allows observation of the renal functional units: the glomeruli, **c**,**d** Tracks of glomeruli. Manual selection, **e**,**f** Tracks from the medulla, **g**,**h** Tracks from the main vessels.

The main regions of the kidney can be easily distinguished through specific track features. In Fig.4, multiple individual tracks were manually chosen and displayed as a function of their velocity and the total distance traveled *D*_*Total*_. All tracks are not on the same scale Four distinct track clusters are evident: green for glomeruli-like tracks, blue for medulla tracks, red for main vessel tracks, and black for outlayer tracks. Glomeruli-like tracks appear slender, forming bundles with spinning microbubbles. Medulla tracks are broader, displaying elongated yet sinuous trajectories. Main vessel tracks, even wider, remain straight. Velocity analysis distinguishes them: medulla tracks move slowly (1-5 mm/s), cortex tracks range from 5 to 10 mm/s, and main vessel tracks accelerate (10-15 mm/s). This observation of cortex velocity aligns with earlier research studies^39,40^. Black outlayer tracks segment into two: high-velocity (12-14 mm/s) tracks cover short distances, possibly due to framerate limitations capturing swift flows. An intermediate-speed outlayer track that might represents a microbubble in the afferent artery, just entering the glomerula or another possible scenario. Remarkably, within the medulla, no track surpasses 0.1 mm in distance owing to its slow pace; the brief acquisition time restricts extended tracking. Lastly, it’s notable that tracks within the medulla and the glomeruli can exhibit comparable speed and travel distance, posing a challenge for differentiation, as discussed further on.

**Figure 4.**
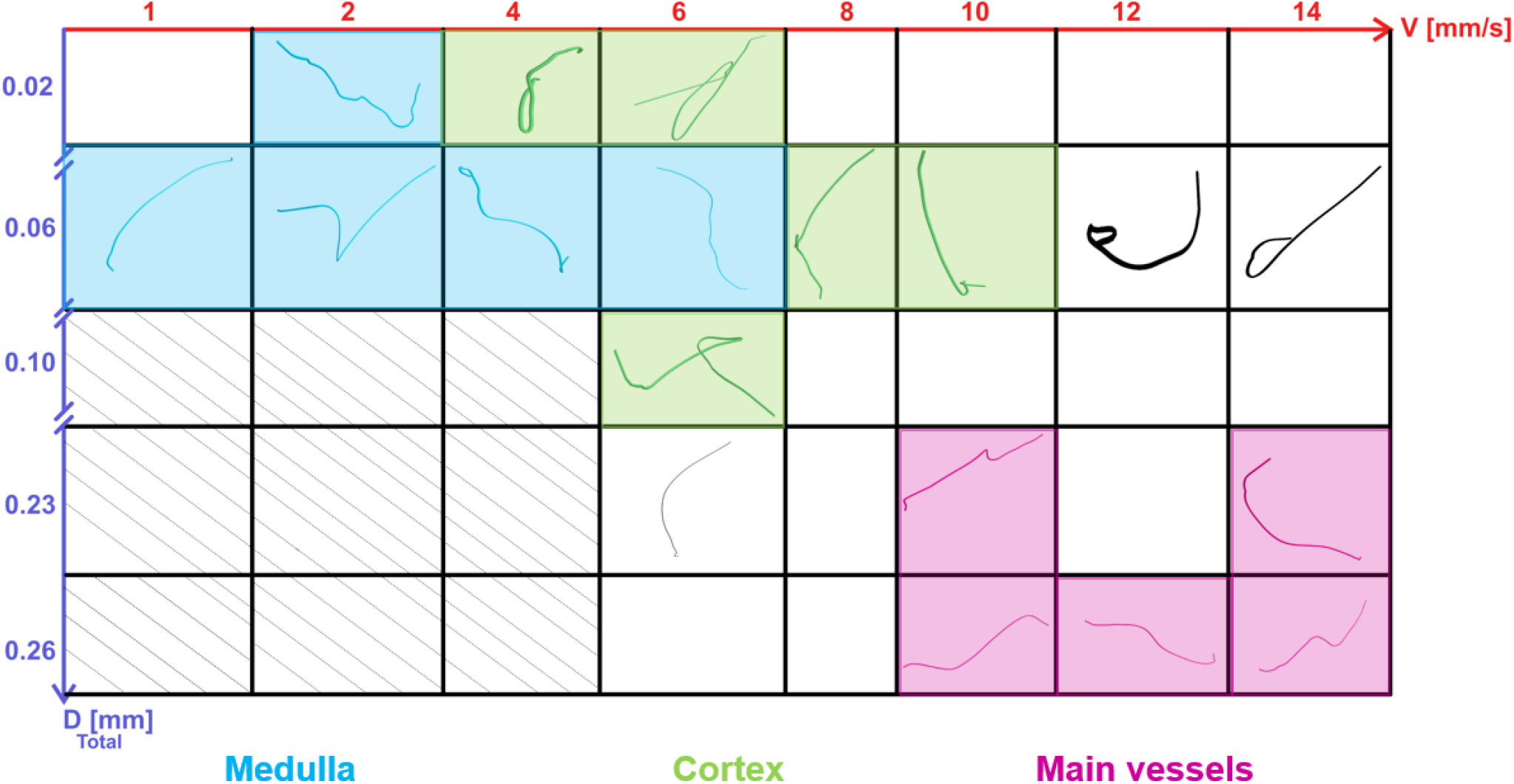
Ethology of 3D sULM tracks: manually selected tracks from all the renial regions. Medullar tracks in light blue. Glomeruli-like tracks in green, Main vessel tracks in red and outlayer tracks in black. All volumetric views are centered on the tracks for displaying clarity, thus tracks do not have the same scale. The tracks are displayed regarding the average velocity on the red x-axis and the total traveled distance ion the green y-axis.

### 1.2 3D metrics to characterise the microcirculation

In order to characterize the kidney’s microcirculation on a macroscopic scale accurately in 3D, we conducted a comparative analysis of metric values across various renal regions in rats. First, volumetric sULM *velocity* rendering is shown in Fig.5 **a**. Values ranging from 1 to 10 mm/s, the low speed corresponds to the tracks in the inner medulla and in the upper region of the cortex, intermediate speeds are more present in the arteries of the cortex and finally high speed dominate in the main vessels (interlobar and arcuate arteries). Two cross sections in the coronal and sagittal planes in **b** and **c** display the velocity values. Statistical analyses of the metrics were represented by boxplot topped by a Student’s statistical test with a confidence level of 95% (Fig.5 **d, f** and **h**) and their spatial representation are displayed in Fig.5 **b, c, e** and **g**.

**Figure 5.**
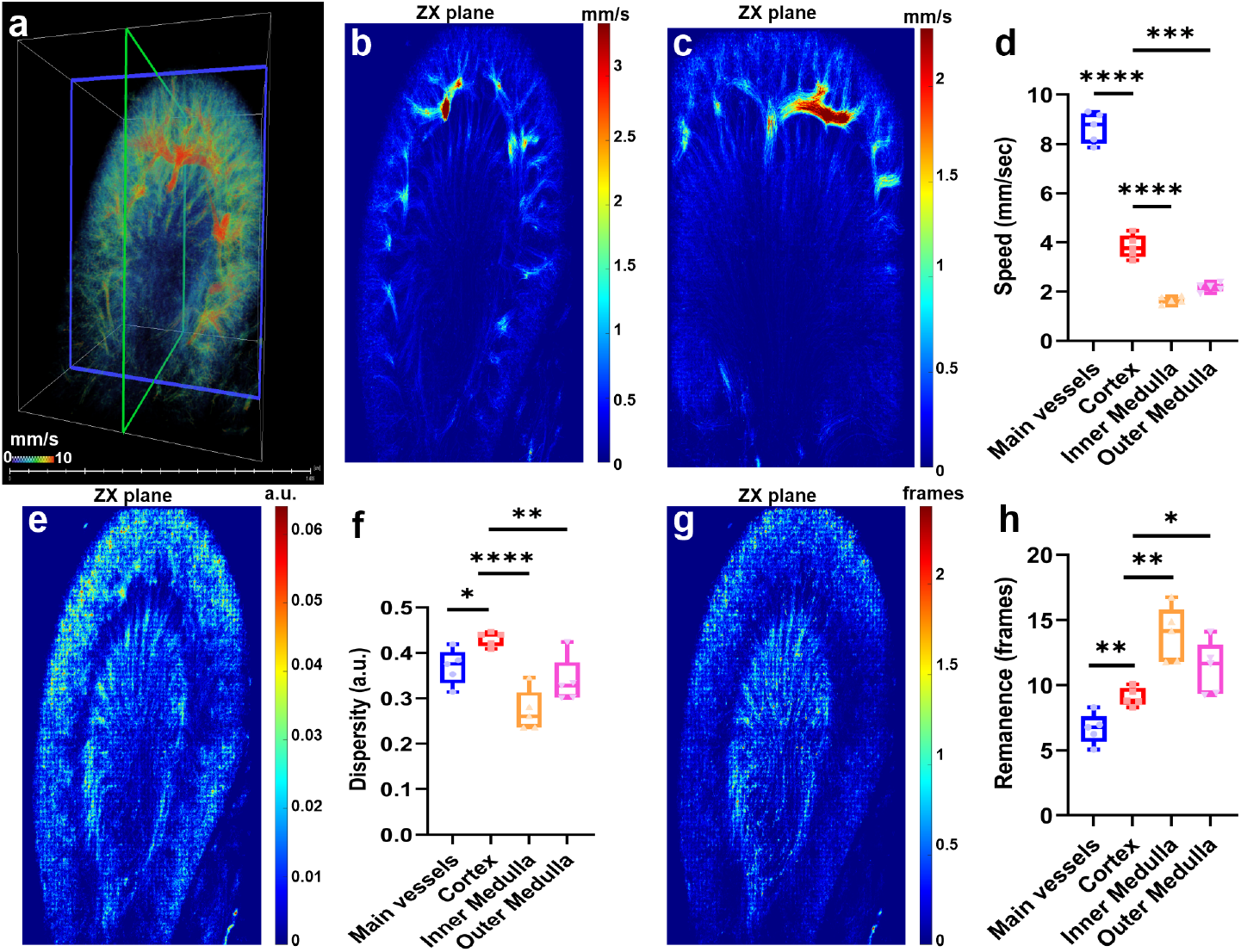
Comparison of the metrics within different renal regions: main vessels, cortex, inner medulla, and outer medulla. **a** Volumetric velocity map. Velocities within the different regions in the coronal (blue) and sagittal (green) planes are shown in **b, c**. Boxplot for *velocity* estimation in **d**. Dispersity within the different regions **e** and its corresponding boxplot **f**. Remanance within the different regions, **g** and its corresponding boxplot **h**.

The boxplot in Fig.5 **d** shows that the *velocity* can be a good discriminater between the four regions of the kidney. The microflow speed is high in the main vessels and almost double the speed in the cortex while the latter is also double the speed in the medulla, the deeper the microbubble flows in the inner medula, the lower their velocity gets. No statistical significance in difference between the inner and outer medulla. Note that the speed in the main vessels should be in order of cm/s in rat’s kidney^19^ hence it is underestimated due to a volumetric imaging rate of 130 frame/s.

The path of the microbubbles found in the medulla, whether internal or external, exhibit statistically slower, more dispersed, and more remanence compared to those present in the cortex (Fig.5 **f** and **h**).

The difference with the values of the metrics between the cortex and the main vessels is statistically significant. For instance, the *remanence* in the cortex is around 10 frames while it is around 6 in the main vessels. Regarding the values of the metrics in the medulla and the glomeruli, they seem to be similar (Fig.4 **f** and **h**). The similarity is hardly surprising, given that both structures are composed of capillaries.

### 1.3 Glomeruli physiological estimation

To estimate physiological parameters like the glomerular filtration rate, it is essential to carefully pinpoint glomeruli within the cortex. This task is achieved through a systematic two-step approach: Firstly, cortical tracks are extracted by applying a velocity threshold (3 *<* track speed *<* 6 mm/s). This velocity parameter serves as a discriminant, effectively distinguishing various segments of the kidney’s structure (refer to Fig. 5 **d**). Subsequently, the Pathway Angular Shift (PAS) metric (outlined in Sec.) is employed on the tracks established via the dual velocity cutoff. If the computed PAS value surpasses a designated threshold of 40 degrees, all data points exhibiting *PAS >* 40° are identified as localizations within the glomerulus. Note that the threshold value used in PAS is completely arbitrary.

Figure 6 depicts the central findings of this study. In Fig.6**a**, 3D sULM rendering of the kidney is displayed, with a closer view provided in **b**. Note that the kidney view in Fig.6 **a** and **b** is the same as Fig.3. Here, glomeruli are highlighted in blue, main vessels in orange, and the medulla in green. In Fig. 6 **c** demonstrates the probability function for the total traveled distance, which reaches its peak at 18 *µ*m. Notably, due to the complex presence of various vessel loops within the capsule, determining the size of the glomeruli through microcirculation is impractical. To approximate a size-related metric, we propose utilizing the total traveled distance within the glomeruli. Upon statistical analysis encompassing all the glomeruli across the rat population, a total traveled distance of 7.50± 0.57 *µ*m is obtained. This measurement is notably smaller than the actual glomerular size (which boasts a 50-micron diameter).

**Figure 6.**
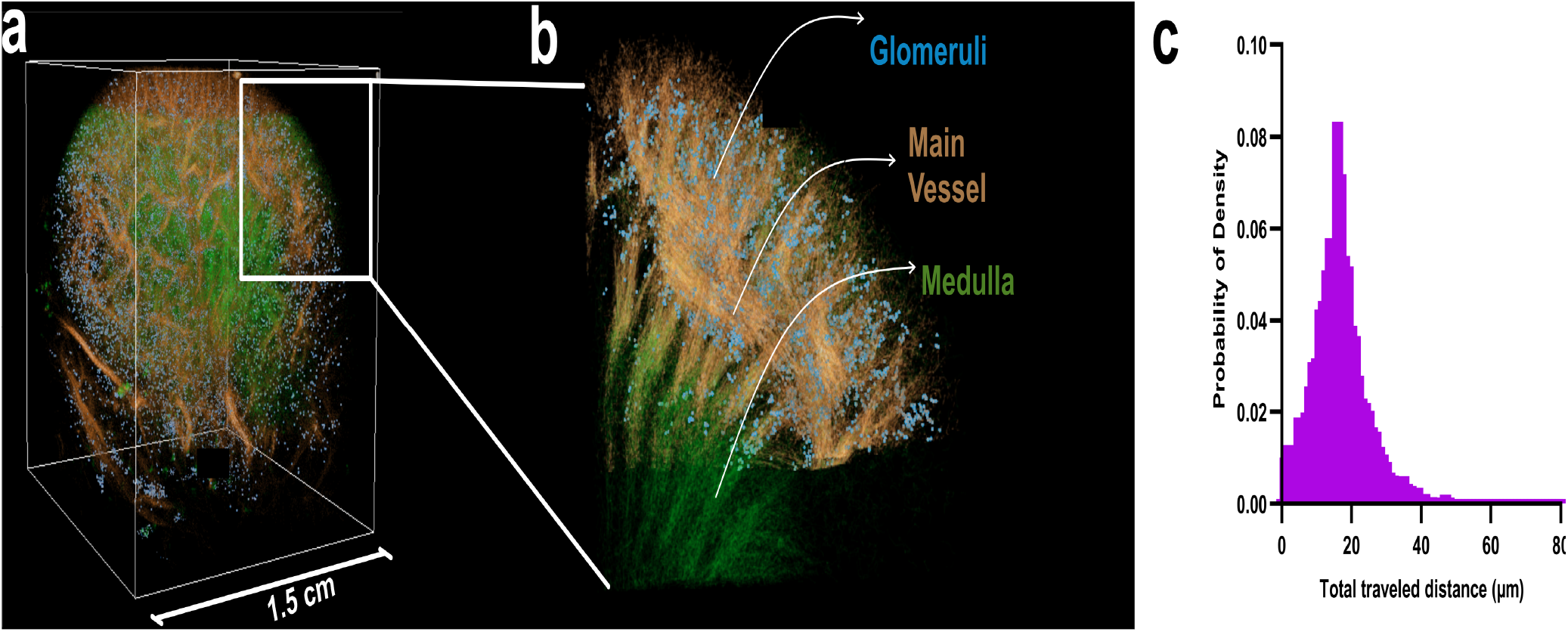
Whole organ volumetric sULM reveals accurate estimation of glomeruli physiology: **a** Volumetric sULM rendering with three different families encoded in color. Medulla in green, main vessels in orange and glomeruli in blue. **b** shows a zoom on a part of the kidney. The probability of density of all the glomeruli tracks (in blue) of the total traveled distance **c**.

**Figure 7.**
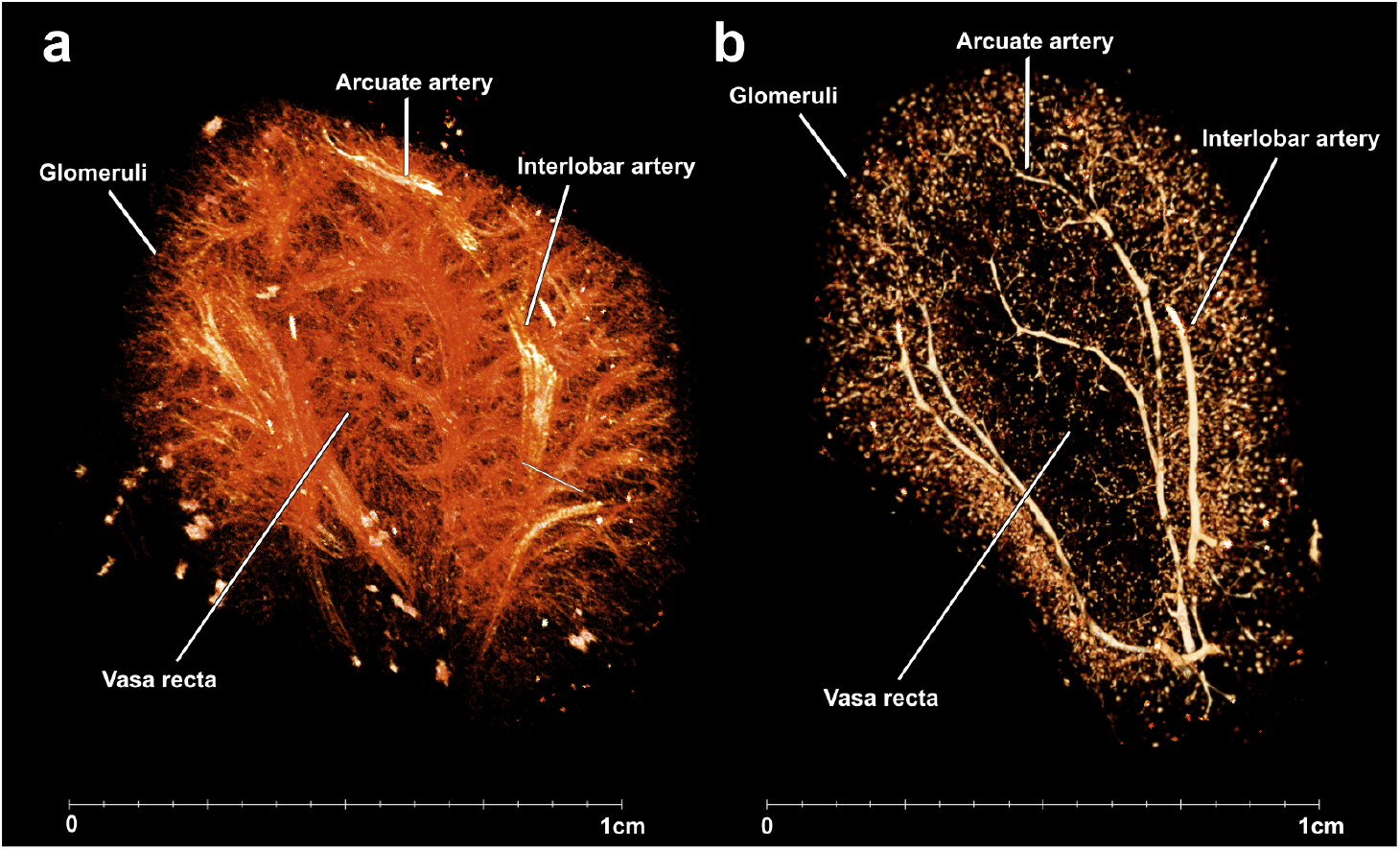
Comparison of the main structures between 3D sULM and micro-CT imaging on the rat kidney. a) 3D sULM, b) Micro-CT.

### 1.4 Ex-vivo reference technique

The preparation of ex vivo specimens for micro-CT imaging is a crucial step that eliminates the influence of neural or chemical signals responsible for vessel constriction or dilation, thus affecting vessel proportions. Nevertheless, it’s vital to acknowledge that the micro-CT imaging process itself can be invasive and may introduce significant artifacts while quantifying results. Factors such as tissue swelling, perfusion pressure during contrast administration, and the effects of contrast curing and paraffin embedding can complicate precise measurements of vessel dimensions. As a qualitative comparison, we observe identical microvascular structures for both techniques as seen in Fig. 7. However, as mentioned in^21^, comparing in-vivo super-resolution ultrasound with ex-vivo microvascular imaging technique (micro-CT) becomes complex. Although micro-CT offers slightly superior resolution (voxel size of 5 *µ*m compared to 10 *µ*m for 3D sULM utilized in this study), 3D sULM presents a more promising alternative due to its non-invasive and in-vivo nature, overcoming suboptimal contrast agent fixation and enabling longitudinal studies with minimal artifacts. Additionally, it’s worth noting that a micro-CT scan can take at least 11 hours, while ULM only requires 5 minutes.

## 2 Conclusions & Limitations

In this study, we proposed a whole organ ultrasound imaging technique. The utilisation of 3D sULM in rat kidneys allowed an accurate volume reconstruction of the microvascular mapping in 5 rats. The high volumetric rate of the acquisition and the low threshold of spatiotemporal filtering made it possible to follow the microbubbles along their intravascular with a single tracking algorithm. In this manner, we achieved a comprehensive examination of microbubble behavior within cappillaries such as in both the vasa recta of the medulla and the nephrons in the cortex. This work allowed us to witness, for the first time, microbubbles circulating within the renal functional units in 3D, in-vivo, in the pre-clinical phase. Additionally, we could observe microbubbles in the medulla, a complex region that remains relatively understudied in nephrology. The vasa recta appears to be correctly reconstructed from the main vessels, and we were able to characterise it using 3D metrics. The use of metrics in each of the renal zones enabled us to compare the behaviour of microbubbles in the different structures of the kidney. Moreover, an innovative glomerular mask was adopted, uniting microflow velocity with the PAS. This straightforward method enables a precise quantitative evaluation of glomerular physiological parameters—an important stride towards GFR estimation.

One of the main limitations of this study lies in the specificity of the metrics used for each of the applications. It will be important to test each metric under various conditions such as: pre-clinical and clinical settings, and evaluating performance across healthy organs and pathologies.

Another limitation stems from the absence of a readily applicable technique for direct comparison with 3D sULM’s high resolution. Micro-CT imaging, the closest candidate, presents challenges due to its invasive nature, potentially causing discomfort to animals and consequently compromising contrast fixation.

Finally, the low-threshold spatiotemporal filtering allowed us to observe very slow flows in the cortex and medulla. Nevertheless, we observed a loss of signal from microbubbles in the center of the vasa recta (medulla). In our opinion, this decrease in intensity is due to the filtering, which considers the microbubble to be quasi-static and coherent (due to its high intensity), and therefore the microbubble signal and the tissue are indistinguishable entities during the process. The lack of precise localization could potentially result in pairing errors within the center of the medulla, creating a”blurred” effect in this region. However, since the medullary region is a relatively understudied capillary complex, this remains a hypothesis that requires further investigation and verification.

### 2.1 Perspectives

The emergence of 3D sULM as a promising technique brings about new possibilities. Its non-invasive, in-vivo characteristics provide distinct advantages. As we delve into optimizing the use of 3D sULM to gain deeper diagnostic insights into various pathologies, it’s important to exercise caution and acknowledge the current limitations it carries. Important validation process involves ensuring its applicability across diverse biological systems and experimental conditions.

From a technological standpoint, the advancement of faster and more powerful hard disks is crucial for enhancing the capabilities of 3D sULM. By having faster data transfer resources, researchers can conduct continuous acquisitions without inter-block waiting times. This dual benefit manifests in two key aspects: Firstly, it extends the visualization duration of microbubbles, effectively transforming them into mobile pressure sensors. These sensors glean insights by dynamically adapting to various environments during their prolonged journeys. The augmented acquisition time empowers these microbubbles to explore diverse conditions, thus yielding a richer trove of information.

Additionally, faster computing capabilities would enable real-time processing and direct visualization of the acquired 3D sULM data. This feature would be invaluable for guiding imaging procedures, allowing researchers to verify the quality of the data as it is being acquired, and potentially making adjustments on-the-fly to optimize the imaging process.

Finally, to ensure accurate localization and tracking of both fast and slow microbubbles, there is a crucial need for more robust clutter filtering techniques. Deep learning-based filters, such as neural networks, offer promising solutions, harnessing their potential to effectively distinguish clutter from microbubble signals. Additionally, exploiting the non-linear properties of contrast agents^23–25^ combined with sULM can further enhance the filtering process.

## Acknowledgment

We thank the”Live Imaging Platform” (Université de Paris-Cité), and specifically Lotfi Slimani for the micro-CT acquisitions with the Skyscan 1172 Bruker.

This study was funded by the European Research Council under the European Union Horizon 586 H2020 program/ERC Consolidator grant agreement No 772786-ResolveStroke.

O.C. holds patents in the field of ultrasound localization microscopy and is a cofounders and shareholders of the Resolve-Stroke startup

